# The Ice Nucleating Protein InaZ is Activated by Low Temperature

**DOI:** 10.1101/2020.05.15.092684

**Authors:** Steven J. Roeters, Thaddeus W. Golbek, Mikkel Bregnhøj, Taner Drace, Sarah Alamdari, Winfried Roseboom, Gertjan Kramer, Tina Šantl-Temkiv, Kai Finster, Sander Woutersen, Jim Pfaendtner, Thomas Boesen, Tobias Weidner

## Abstract

Ice-nucleation active (INA) bacteria can promote the growth of ice more effectively than any other known material. Utilizing specialized ice-nucleating proteins (INPros), they obtain nutrients from plants by inducing frost damage and, when airborne in the atmosphere, they drive ice nucleation within clouds and may affect global precipitation patterns. Despite their evident environmental importance, the molecular mechanisms behind INPro-induced freezing have remained largely elusive. In the present study, we investigated the folding and the structural basis for interactions between water and the ice-nucleating protein InaZ from the INA bacterium *Pseudomonas syringae* strain R10.79. Using vibrational sum-frequency generation and two-dimensional infrared spectroscopy, we demonstrate that the ice-active repeats of InaZ adopt a β-helical structure in solution and at water surfaces. In this configuration, hydrogen bonding between INPros and water molecules imposes structural ordering on the adjacent water network. The observed order of water increases as the interface is cooled to temperatures close to the melting point of water. Experimental SFG data combined with spectral calculations and molecular-dynamics simulations shows that the INPro reorients at lower temperatures. We suggest that the reorientation can enhance order-inducing water interactions and, thereby, the effectiveness of ice nucleation by InaZ.

## 1. Introduction

Ice-nucleation active (INA) bacteria are the most effective ice nucleators known.^5, 6, 7^ Using surface-exposed, ice-nucleating proteins (INPros) attached to the outer membrane, they promote ice growth at high sub-zero temperatures.^8^ Water and ice are fundamental for life on Earth in general and atmospheric processes in particular. In this context, INA bacteria play important roles as they affect surface water, the hydrological cycle, local and global climate, and vegetation.^9, 10, 11^ Recent field studies have discovered large amounts of INA bacteria at high altitudes.^12^ In the atmosphere, bacteria have been identified as an important cause of cloud glaciation, snow and hail precipitation, and cloud formation.^9, 10, 13^ Pure water droplets in the atmosphere that are getting super-cooled can remain liquid down to approximately −40°C. It has been shown that, in the presence of *P. syringae cells*, water freezes at temperatures as high as −2°C.^5, 14^ Due to that property, INA bacteria may play an important role especially for mixed-phase, low-altitude clouds, where abiotic particles are far less effective ice nucleating particles than bacterial cells.^5^

**Scheme 1.**
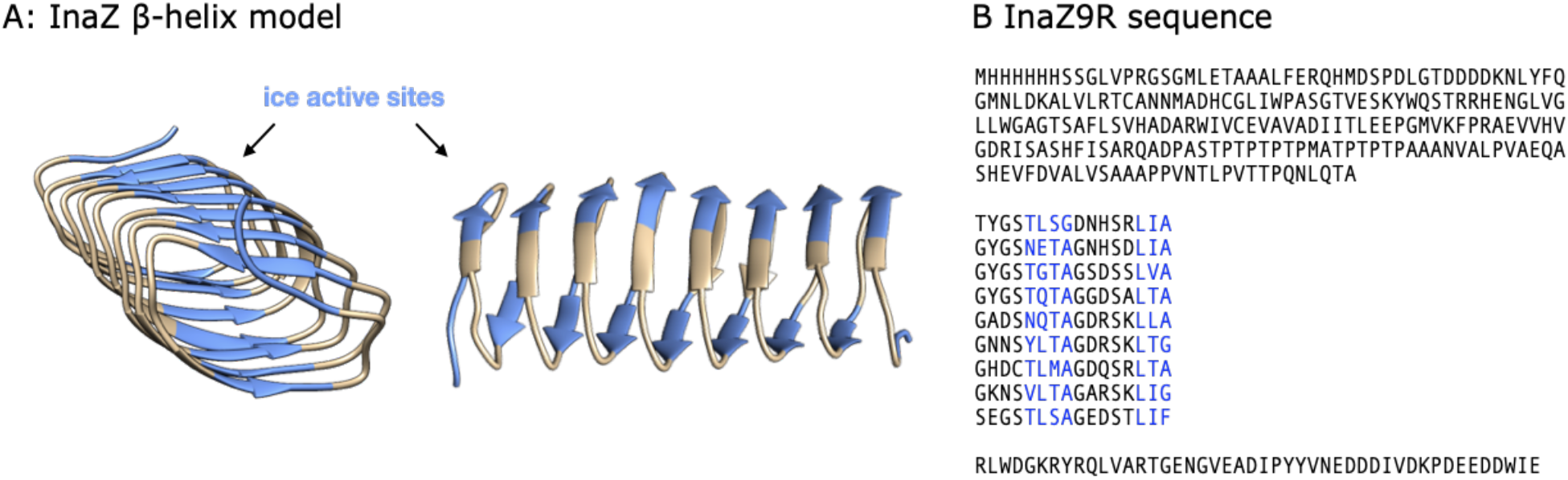
Illustration of the InaZ repeat sequence and the model inaZ9R construct. Based on current models, the blue sequences are ice-nucleation active. (A) Rendering of a part of the repeat structure of the ice-nucleation active protein InaZ. (B) Sequence of the InaZ9R construct. It contains an N-terminal tag, the membrane binding N-terminal domain, the ice-binding repeat units, and a C-terminal domain of unknown function.^1, 2, 3, 4^

Due to this broad impact, INA bacteria and their capability to control water’s aggregation state may be an integral part of sustaining Earth’s ecosystems.^15^

INA bacteria also have significant economic implications since they cause severe frost damage on crops and other types of vegetation. At the same time, *P. syringae* has also caught attention for technological applications such as artificial snow formation, food preservation, cryo-medicine and freezing technologies.^6, 16, 17, 18^ However, true rational design of artificial biomimetic icing materials is still impeded by the lack of a molecular picture of how INPros catalyze ice nucleation.

Despite the numerous environmental and economic aspects, where bacterial ice nucleation is of critical importance,^8, 11^ the fundamental mechanisms of protein-driven ice nucleation are still not clear. While the InaZ sequence containing ^~^1200 amino acid residues has been known for some time,^4, 19^ and there have been several theoretical studies of INPro folding,^3, 20, 21, 22^ only limited experimental evidence on the structure of INPros is available.^19, 23, 24, 25, 26, 27, 28^ Previous vibrational sum-frequency generation (SFG) studies with cultures of *P. syringae* have shown that InaZ can order water and promote energy transfer across the water interface.^29, 30^ It has also become evident that the ability of *P. syringae* InaZ to control water orientation is enhanced when cooled to temperatures close to the water melting point.^29, 31^ However, since bacterial cell surfaces contain a variety of many different proteins and other biomolecules, these studies could not track the molecular origin of this activation at the protein level. Our current picture of the interaction of INPros with water is still, to a large extent, based on theoretical models and molecular dynamics (MD) simulations^3, 32, 33^, with contradicting conclusions for protein folding and the role of side chain motifs. The first model was put forward by Kajava and Lindow and is based on parallel β-sheets.^21^ Later, based on homology modeling, Graether and coworkers proposed that InaZ folds into a β-helical structure, similar to insect antifreeze proteins.^3, 32, 34^ In the latter, more current picture based on MD simulations, INPros bind ice through highly conserved repeat units in a fashion that is similar to some antifreeze proteins (Scheme 1). The structural similarity between INPro and antifreeze proteins has been proposed due to the similarity in the ice-binding sequence (TxT motifs) and is supported by experimental reports that these proteins have overlapping functions of ice-nucleation and ice-growth-inhibition,^24, 35^ which are proposed to depend on the protein size. INPro are larger and therefore more efficient in nucleating ice and promote freezing at higher subzero temperatures compared to smaller antifreeze proteins. Size also affects the ice nucleation efficiency of INPro, which have been shown by experimental studies to oligomerize into larger units that promote ice formation at higher temperatures than protein monomers.^28, 36, 37^ Studies on INA bacteria suggest that lowering the temperature induces the ice-nucleation activity of INPros through reorganization and concentration of INPros at cellular poles.^38^

The current lack of experimental data about INPro activity can be traced back to the general challenges involved in studying the conformation of proteins *in situ* at interfaces. Here, we have used SFG spectroscopy to probe the structure of the ice nucleating protein InaZ at the air-water interface, as a model system for the cell-water interface. SFG relies on the resonant enhancement of frequency mixing between visible and infrared laser pulses, when the infrared pulse is resonant with a surface vibration. The selection rules of SFG dictate that only ordered species at an interface are detected. Therefore, in the amide-I region, SFG can probe the structure of interfacial proteins and in the O–H stretching region the technique can observe water molecules interacting with the protein interface. In the present study, we report on experiments with isolated model INPros and elucidate the activation mechanism. We used a truncated version of the InaZ that contains 9 out of 67 repeat units (repeats 1-4 and 63-67) of the native ice-nucleation active protein (see Scheme 1) of *P. syringae* strain R10.79. Using this construct (termed InaZ9R), we address the question whether the ordering of water molecules previously observed for INA bacteria is associated with INPro molecules. We provide experimental evidence that the central repetitive region of InaZ acquires a β-helical structure as proposed by recent modeling studies, and that InaZ reorients at low temperatures to increase contact to water molecules and, thereby, ice-nucleation activity.

## 2. Results and Discussion

### 2.1 Water Structure at the InaZ Interface

Using SFG spectroscopy in the water region of the vibrational spectrum, we have monitored the interaction of water in direct contact with InaZ9R. Figure 1 displays temperature dependent SFG spectra collected in ssp (s-polarized SFG, s-polarized visible, and p-polarized infrared) polarization for InaZ9R in PBS buffer at the air–water interface compared to neat PBS buffer spectra. For technical reasons, heavy water was used to prepare the buffer solutions (see SI). The spectra of InaZ9R at the air–water interface show resonances near 2363, 2408, and 2485 cm^−1^ related to strongly, intermediately strong and weakly hydrogen bonded water molecules at biological surfaces, respectively. There is also a weak, but discernible signature of the free O–D resonance near 2655 cm^−1^. Already at 20°C, the intensities of the observed resonances are significantly stronger when InaZ9R is present, compared with the neat buffer surface spectrum shown in panel B, indicating an increased water order. This is often observed for biological surfaces such as protein films and lipid monolayers, and can be explained by the interaction and alignment of water molecules at the InaZ9R interface.^39^ As the buffer temperature is decreased to 10°C, and subsequently to 5°C (close to the melting temperature of heavy water of ^~^4°C), the water signal is strongly increased. Fitting the data with Lorentzian functions for the resonant modes and a non-resonant background (see SI for details) shows that all water modes are affected by the temperature change, to slightly varying extents (see Figure 1C).

**Figure 1.**
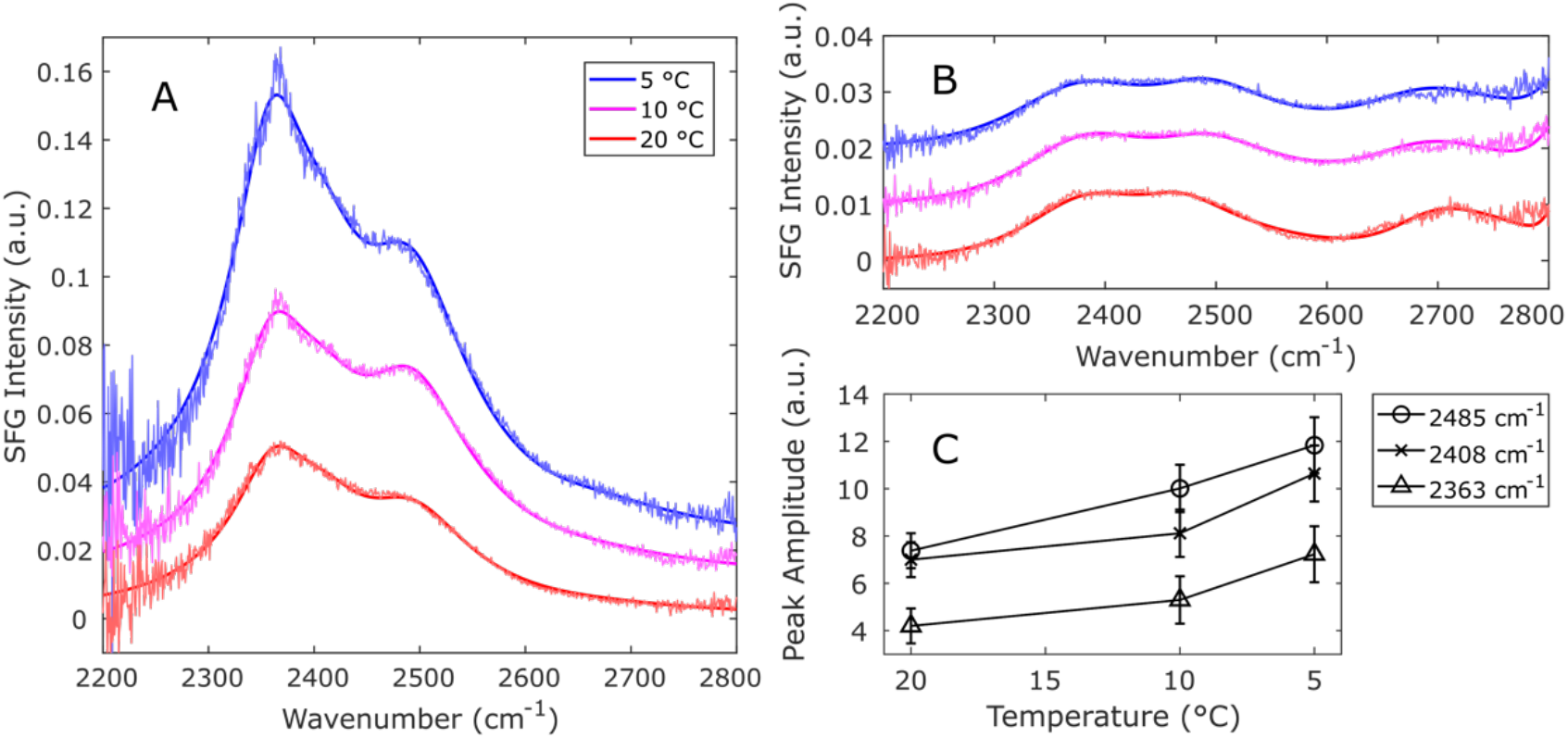
Temperature-dependent OD-stretch SFG spectra collected in the ssp polarization combination, along with fits. The spectra are slightly offset for clarity. (A) Spectra of water molecules interacting with the ice-nucleating active protein InaZ9R in D_2_O-based PBS buffer. The intensity of the water signal is increased significantly for lower temperatures in the presence of inaZ9R. (B) Spectra of the neat buffer surfaces remain largely unchanged. (C) Plot of the SFG amplitudes for the different water modes fitted to the InaZ9R spectra.

At the same time, SFG spectra of the neat PBS buffer surface (Figure 1B) do not change appreciably with temperature. This is in agreement with previous studies of non-ice nucleating interfaces.^29, 40^ In those reports, an increase in the water signal has only been observed for ice-active antifreeze proteins and ice-nucleating bacteria. For non-ice nucleating (bio)surfaces, such as the air–water interface, lipids, sugars and non-ice active proteins or bacteria, it has been shown that the water signal is not affected by such a temperature decrease. Clearly, the INPro InaZ9R is ordering water effectively and its control of the water structure becomes more efficient at temperatures near its ‘operating temperature’; temperatures close to the water melting point at which INPros are typically active. The similarity between these temperature-dependent InaZ spectra and previously published temperature-dependent *P. syringae* spectra, provides strong experimental evidence that InaZ is indeed the water-ordering agent responsible for the ice-nucleating activity of these bacterial cells.

### 2.2 InaZ Secondary Structure in Solution

This raises the question as to the structural basis for the enhanced water order at lower temperatures. Previous studies with INA bacterial cells have not been able to answer this question because of the structural heterogeneity of bacterial cell surfaces. With the purified InaZ9R construct, we now attempt to address this question – by firstly determining the solution-state structure of InaZ9R and thereafter tracking the binding geometry of InaZ9R at the air–water interface.

To determine the secondary structure in solution, we have used conventional Fourier-transform infrared (FT-IR), two-dimensional infrared (2D-IR) and UV circular-dichroism (UV-CD) spectroscopy. Figure 2A displays the experimental FT-IR and 2D-IR amide-I (1600-1700 cm^−1^) spectra of InaZ9R in PBS buffer. 2D-IR Spectra are recorded by exciting the protein solution at a specific frequency ***ω***_pump_ and probing the resulting changes in the absorption over the entire IR frequency range. Scanning ***ω***_pump_ and plotting the absorption change as a function of ***ω***_pump_ and ***ω***_probe_, we obtain two-dimensional IR spectra that can be regarded as the vibrational analog of 2D-NMR spectra.^41^ When two normal modes A and B are coupled, exciting mode A (***ω***_pump_ = ***ω***_A_) causes an absorption change at the frequency of mode B (***ω***_probe_ = ***ω***_B_) and vice versa, and this gives rise to cross-peak patterns at (***ω***_A_, ***ω***_B_) and (***ω***_B_, ***ω***_A_) in the 2D-IR spectrum. In the case of the amide-I mode of proteins the individual cross peaks can generally not be resolved, but the shape of the amide-I 2D-IR spectrum is still very sensitive to the secondary structure.^42^ The FT- and 2D-IR spectra are composed of a single broad band that peaks at 1632 cm^−1^, and the 2D-IR spectrum does not contain any strong cross peak. The spectra show a close resemblance to previously measured IR spectra of proteins composed mainly of β-helices, with segments of other secondary structure.^43^ To test which of the theoretical models agrees best with the experimental data, we have calculated 2D-IR spectra based on both the β-sheet and the β-helix model to compare with the experimental data (Figure 2B and C). For the β-sheet model, we based the calculation on the published 48-mer repeat unit composed of stacked antiparallel β-sheets.^21^ For the β-helix model, we used a structural model that was proposed in a previous INPro simulation study.^3^

**Figure 2.**
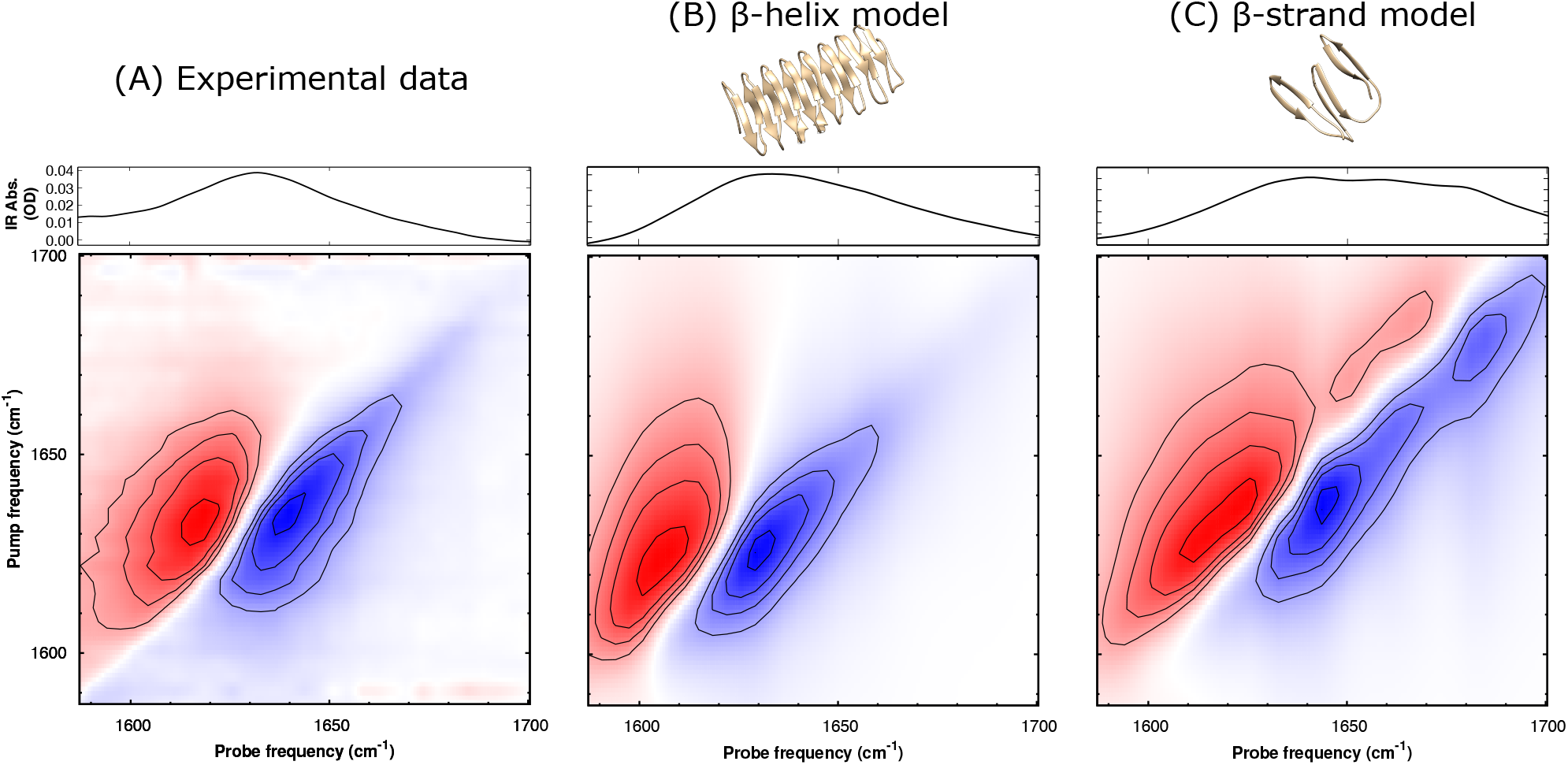
Experimental and calculated FT-IR (top) and parallel 2D-IR (bottom) spectra of InaZ9R in solution. The 2D-IR spectrum is sensitive to the secondary structure of a protein^41^ and can be regarded as a fingerprint of its conformation. Comparison of experimental and calculated spectra shows that the β-helix model captures the experimental data very well, while the β-sheet model shows significant deviations between theory and experiment.

The models were briefly equilibrated using a 100 ns GROMACS simulation (see SI for details). As the stacked antiparallel β-sheet model was unstable over the course of the simulation, we artificially stabilized the structure by introducing distance constraints along the aligned β-sheets. The 2D-IR spectra were calculated using an amide-I Hamiltonian model applied to this restrained simulation for the β-sheet hypothesis, while the spectral calculations for the β-helix model were performed on an unrestrained simulation with otherwise similar conditions (see SI for details). To calculate the spectra depicted in Figure 2B and C, an additional inhomogeneous broadening was applied to the local-mode frequencies in order to match the broad experimental peak shape. This is probably due to the C- and N-termini of the full InaZ9R sequence used in the experiments (see scheme 1) that are likely to adopt a less-well defined, (partly) random-coil structure in solution, as is also indicated by UV-CD spectroscopy (figure S4). We used the same inhomogeneous bandwidth for both models to ensure a fair comparison. Both with and without the applied inhomogeneous broadening, it is clear that the β-helical model results in a closer spectral match than the β-sheet model (see figure S5 for comparison without inhomogeneous broadening). The calculated β-sheet model spectra are significantly shifted with respect to the experimental resonance positions, and exhibit high-frequency modes that are absent in the experimental spectra. The 2D-IR spectra calculated for the β-helix model, on the other hand, capture the experimental spectra very well. Minor deviations of experiment and calculation observed for the β-helix model might be explained by spectral contributions of coiled structures in the N- and C-terminal domains that are different from the purely random-coil structure (here modelled by the included inhomogeneous broadening) and the oligomerization of InaZ9R. Oligomerization has been reported previously for other InaZ constructs and related β-helix structures of hyperactive insect anti-freeze proteins (AFPs).^19, 27, 44, 45^ In summary, we conclude that the β-helix model agrees with the IR data and describes the solution structure of InaZ9R well.

### 2.3 InaZ Activation at Low Temperatures

To explain how the activity of InaZ is promoted at temperatures close to the water melting point, we have recorded amide-I SFG spectra of InaZ9R at the air–water interface. Amide-I SFG spectra are, like amide-I IR spectra, sensitive to secondary structure, and because it is a coherence-based technique,^46^ it can also provide information about the orientation of interfacial proteins. Figure 3 displays amide-I SFG spectra collected at room temperature (20°C) and close to the water melting temperature (5°C). The spectra recorded at 20°C show a mode centered near 1640 cm^−1^ and a second mode near 1720 cm^−1^. The 1720 cm^−1^ mode can be assigned to lipid molecules, which indicates the presence of protein-associated lipids. Both SDS-PAGE and intact protein mass spectrometry point to an increased molecular weight of the protein compared to the expected mass based on its assumed amino acid sequence (see SI). The feature near 1640 cm^−1^ can be assigned to β-sheet type protein structures.^47, 48, 49^ Since SFG selection rules dictate that only ordered surface species are detected, the presence of the amide-I mode clearly shows that InaZ9R forms an ordered layer of proteins at the air-water interface, which is induced by low temperature exposure. This is also supported by SFG spectra in the C–H stretching region, where aliphatic and aromatic modes are visible and demonstrate a high degree of alignment within the hydrophobic side chains (see SI for data).

**Figure 3.**
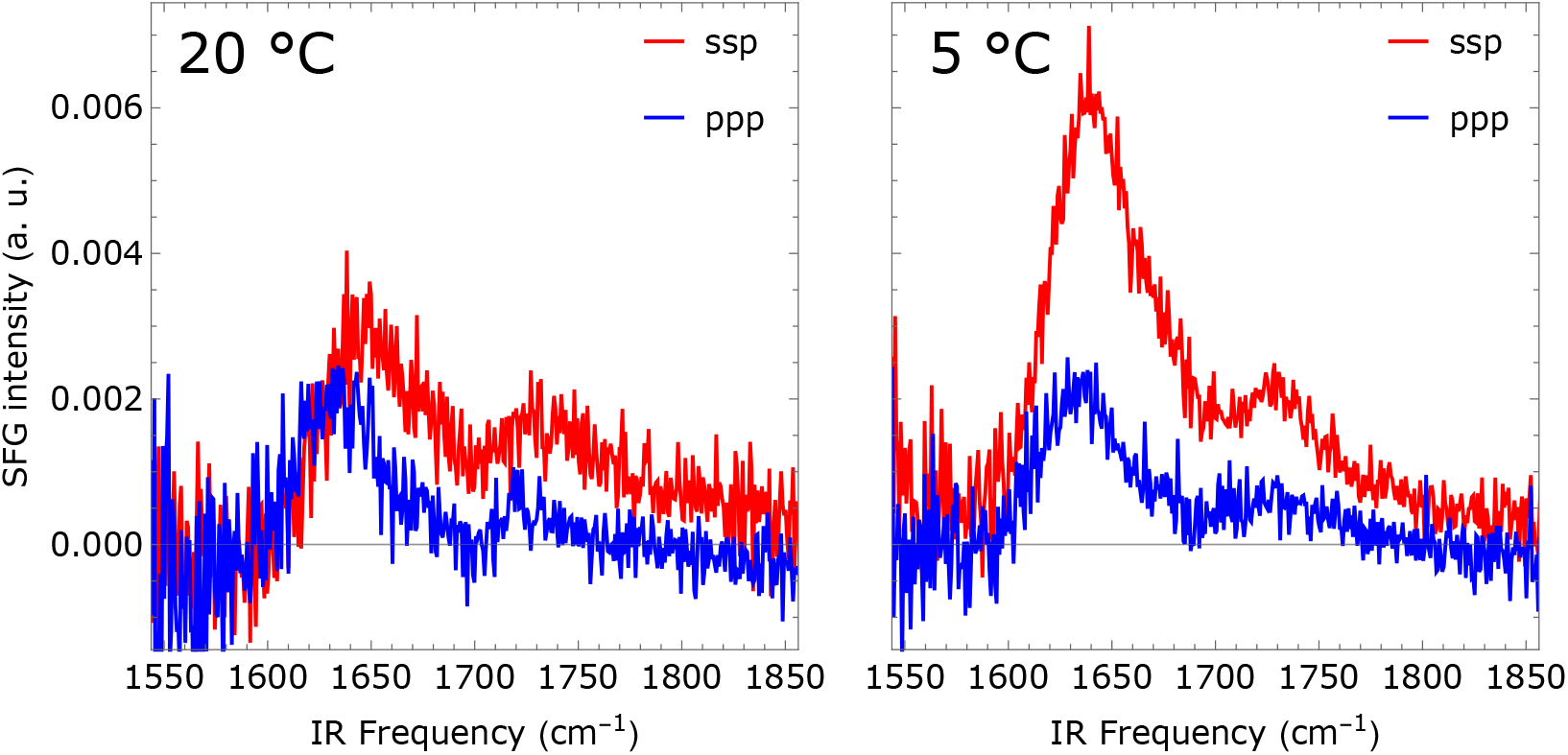
Temperature-dependent amide-I SFG spectra of InaZ9R at the air-water interface in ssp and ppp polarization. (A) Spectra recorded at 20°C show a resonance near 1640 cm^−1^ related to protein β-sheets and a resonance near 1720 cm^−1^ related to lipids. (B) Spectra collected for 5°C show a strong increase in amide-I ssp intensity which indicates the protein has changed orientation at the interface.

At 5°C, the resonance positions within the SFG spectra remain largely unchanged while the peak ratio of the 1640 cm^−1^ protein resonance between the ssp and ppp polarization combinations is dramatically increased. The fact that the resonance positions are similar at both temperatures indicates that the folding of InaZ9R does not change considerably.

Often, a change in amide-I peak ratios indicates a reorientation within protein monolayers.^50, 51, 52^ The Chen group has developed methods to determine the orientation of helix and β-sheet structures.^53, 54^ However, to the best of our knowledge, the SFG response of a β-helix, has never been analyzed or quantified before with SFG. Since the SFG response of InaZ9R can for this reason not be determined by fitting with Lorentzian functions or by spectral inspection, we have calculated SFG spectra based on the β-helix model, which encompasses the central repeat domain of InaZ^3^. While it is known that InaZ is membrane anchored with the N-terminal domain,^1, 2, 3^ the orientation of the protein with respect to the cell surface is not known. Figure 4A presents an overview of the match calculated and experimental SFG spectra for a range of combinations of tilt and twist angles (θ, Ψ)^55^ with a step size of 2.5°, indicated by the residual sum-of-squares (RSS). There is a narrow range of angles for which theory and experiment closely match. The smallest RSS, and thus the closest spectral match, is found to be for small tilt angles for 20°C, while the minimal RSS values for 5°C are found at larger tilt angles. Going from 20 to 5°C, the closest match between the experimental and calculated spectra indicates a transition from (θ, Ψ) = (21°,45°) to (59°, 290°) (see fig. 4B). The calculated spectra capture both the resonance positions and the relative intensities of the experimental data well. This orientational transition would result in an increased water-exposure of the ice-nucleation active sites, through an increased tilt and twist angle (fig. 4C). Specifically, at 20°C, the best-fitting protein orientation indicates a near-parallel alignment of the helix axis to the water surface, with little solvent exposure of the INA sites. At 5°C, the orientation of the INA sites is changed in the best matching orientation, with a larger tilt angle θ and a Ψ rotation around the helix’ axis. In conclusion, the observed cooling-induced change in the intensity ratio between the different SFG polarization combinations can be explained by a protein reorientation that results in an increased solvent exposure of the ice-nucleation active sites.

**Figure 4.**
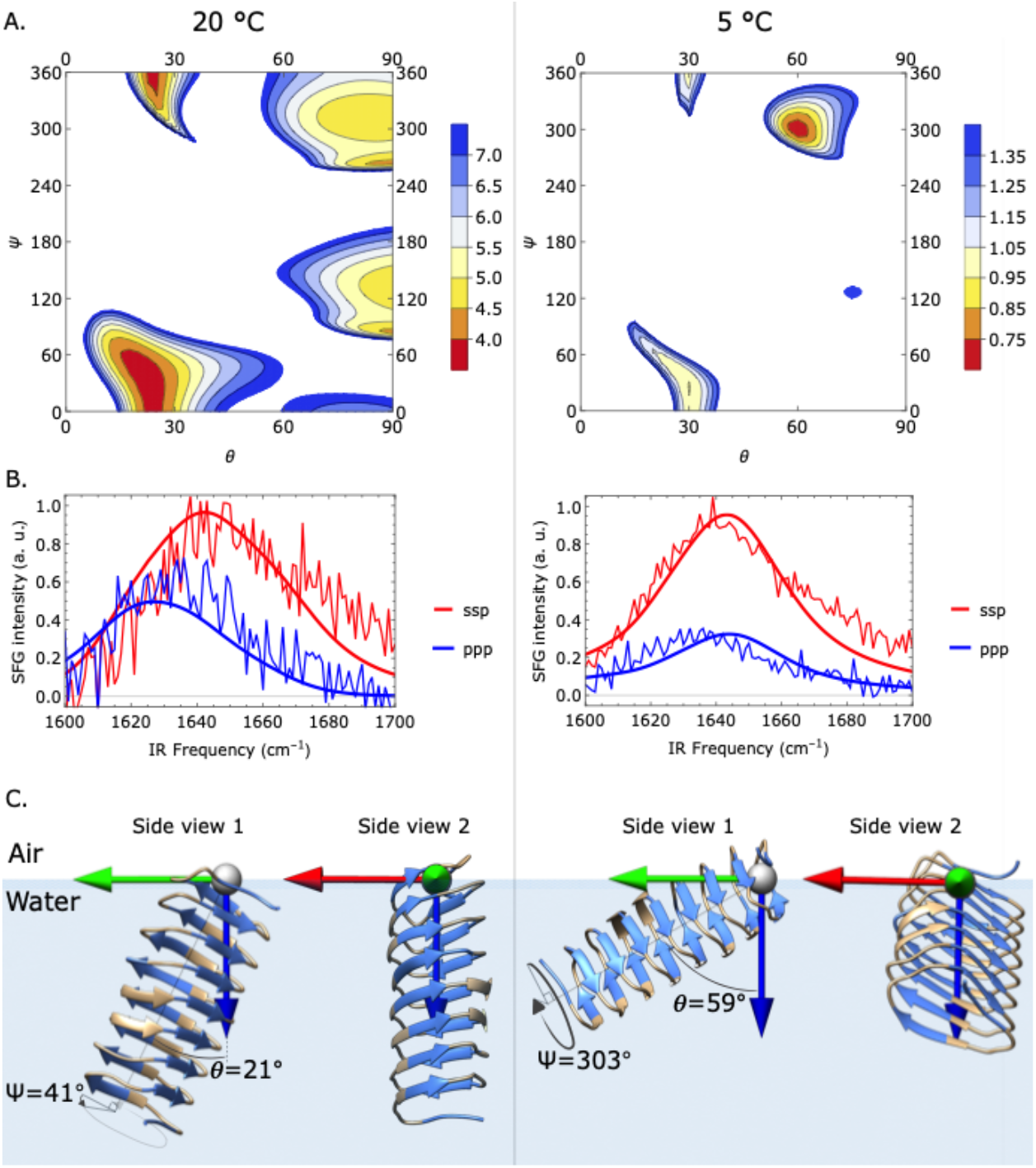
Theoretical and experimental SFG spectra in the amide-I region for InaZ9R at the air-water interface and inferred surface model for InaZ9R based from the calculated SFG amide-I spectra. (A) Residual sum-of-squares (RSS) plots illustrating the spectral match between theory and experiment for different tilt (ϑ) and twist (Ψ) angles. The redder the contour region is colored, the closer the match between the calculated and experimental spectra. The white regions indicate (ϑ,Ψ) values for which the RSS values are more than twice as large as the minimal RSS value. (B) Calculated spectra resulting from (ϑ,Ψ) fits starting at the minimum RSS position, resulting in best matching tilt and twist angles of ϑ=21° ^+31°^_-40°_ and Ψ=41° ^+72°^_-110°_ for 20°C, and ϑ=59° ^+18°^_-8°_ and Ψ =303° ^+42°^_-36°_ for 5°C (positive and negative uncertainties are given as super- and subscripts, respectively, as determined by performing fits with the respective angle fixed at values deviating from its optimal value, until the RSS is twice as high as at the optimal (ϑ,Ψ) position, see SI). (C) Schematic representation of the best matching protein orientations. InaZ9R reorients at lower temperatures with a more inclined orientation at 5°C. In a tightly packed protein layer, the low-temperature pose increases the interaction of the INA site (marked blue) with interfacial water.

## 3. Conclusion

The IR data provides experimental evidence that the central repeat region of InaZ has a β-helical fold. In addition, the SFG data proofs that this pose is also stable at the air–water interface, a simplified model system for hydrophobic surfaces such as cell surfaces. Simulations have been predicting a β-helical folding motive based on sequence homology with insect AFPs.^3, 34^ This study now provides evidence that indeed bacterial INPs and insect AFPs have a similar β-helical structure, despite having contrary effects on water molecules. The difference in ice activity is probably caused by the difference in size of the ice-interacting surface – ice nucleating proteins are five to ten times larger than insect AFPs.

Earlier studies have reported that the ice activity of *P. syringae* is increased when the bacteria are exposed to temperatures around 4°C.^38^ The SFG analysis of the interfacial protein structure shows that this effect may be driven by InaZ9R reorientation when cooled to lower temperatures. Figure 4C provides an overview of the temperature-induced transitions. While at room temperature the long axis of InaZ9R is oriented perpendicular to the surface, at lower temperatures the protein reorients and adopts a configuration that is orientated more parallel to the interface. In addition, the inaZ9R rotates along the long helix axis such that the ice-active strands are oriented more parallel to the surface. The low-temperature geometry increases the exposure of ice-active sites to the interfacial water network when the protein is part of a densely packed layer, which can be expected to form under the experimental conditions applied in this study. Lateral assembly of InaZ in the room-temperature state will lead to burial of ice-nucleating active sites within the protein film, while at low temperatures the protein film exposes a large area of INA sites to the water. This enhances water ordering and is thereby the probable underlying cause of the observed increase in water signal at low temperatures.^56^ Large areas of ice-nucleation active sites are crucial for heterogeneous ice nucleation.^44, 56^ At cell surfaces, based on current models, InaZ is anchored through its N-terminal domain^3^ and would presumably be flexible enough to reorient from perpendicular to parallel to the cell surface with decreasing temperatures. Thus, a reorientation of InaZ could also explain the puzzle of why water close to ice-active bacterial cells becomes more ordered when cooled to lower temperatures.

The cause for the reorientation of InaZ9R is best explained by a subtle shift in the sensitive balance of forces between protein side chains, water molecules and lateral protein interactions. Studies of INpros and related AFP β-helices have shown that all these factors can be affected by temperature changes. For example, it has been shown that hydration^57, 58^ and lateral protein interactions^58^ change significantly when lowering the temperature close to the water melting point. The protein-water interface has also been observed to become less dynamic at lower temperatures.^33^ Together, these factors can surely alter interaction sites for protein-protein as well as protein-water contacts and impact the lateral organization on InaZ at interfaces.

In summary, the presented data provides direct experimental evidence that InaZ adopts a β-helical fold in solution and at the water interface. Furthermore, the data proofs that InaZ is indeed the driving force for the water ordering at lower temperatures observed for *P. syringae* cells. The analysis shows, that InaZ reorients at the interface when it is cooled from room temperature to the ‘operating temperature’ of ice-nucleating proteins, leading to an increased exposure of the ice-nucleation active sites. This temperature-induced activation leads to an increased water order and, consequently, surface freezing.

## 4. Methods and Materials

Details of all methods, materials, and calculations are given in the SI.

### Protein purification

The truncated InaZ protein construct (termed InaZ9R), containing 9 repeat units, was prepared as described previously for a 16 repeat InaZ construct^37^. The InaZ9R sequence was cloned into the pET30 Ek/LIC vector (Novagen, Merck Biosciences) according to manufacturer’s instructions, and used for transformation of *E. coli* Rosetta (DE3) competent cells. A large-scale culture was grown in LB medium and induced with 1mM IPTG over night at 20°C. The protein was purified by immobilized metal affinity chromatography. Fractions containing InaZ9R were subsequently loaded on an anion exchange chromatography column attached to a FPLC system and finally on a size exclusion chromatography column attached to a FPLC system. The individual fractions were either flash frozen in liquid nitrogen and stored for later use in the size exclusion buffer (20 mM Tris-HCl pH 7.5; 150 mM NaCl), or buffer-exchanged into (D_2_O) PBS buffer for further characterization.

### Vibrational spectroscopy: *2D-IR*

The setup and procedures used to record and process 2D-IR spectra have previously been described^59^. Briefly, transmission 2D-IR spectra were recorded by overlapping a chopped narrowband pump IR laser pulse and a broadband probe IR laser pulse on the sample with a delay of 1.5 ps. An additional reference beam, identical to the probe, was passed through the sample at a few mm displacement. The difference-absorption spectrum between the unchopped and chopped pulses were recorded on a spectrograph with an IR-sensitive MCT (HgCdTe) array, and corrected for peak-to-peak stability with the reference beam. The 7 μL IR samples (prepared in D_2_O buffer, to prevent overlap with the OH-bending mode) were pipetted in between two CaF2 windows that were spaced 50 μm and sealed by a greased Teflon spacer.

### Vibrational spectroscopy: *SFG*

Likewise, the setup and procedures used to record SFG spectra have been described before^60^. SFG spectra were recorded in standard reflection geometry, by overlapping a tunable broadband IR laser pulse with a narrowband visible laser pulse in space and time. The generated SFG light was routed to a spectrograph and recorded with an EMCCD camera. All spectra were background subtracted and normalized by a spectrum from a neat gold surface. Protein samples were prepared in phosphate buffered D2O, to avoid spectral interference from water bending modes.

### Spectral calculations: *2D-IR*

A two-exciton amide-I Hamiltonian was constructed based on the solution state MD trajectory at room temperature. The couplings in the Hamiltonian were based on the transition-dipole coupling model^61^ for non-nearest neighbors, whose interaction is dominated by through-space effects, while the nearest neighbor couplings, dominated by through-bond effects, are modelled with a parameterized map of an *ab initio* calculation with the 6-31G+(d) basis set and B3LYP-functional^62^. The local-mode frequencies on the diagonal of the Hamiltonian are estimated using the same hydrogenbond-shift model as published previously^63^, which is based on the effect that various types of hydrogenbonds to the amide group have on the amide-I frequency^64^. By diagonalizing the Hamiltonian we obtain the amide-I eigenmodes and eigenvalues from which the 2D-IR spectra are calculated for each of the frames, after which the response is averaged over all frames.

### Spectral calculations: *SFG*

A one-exciton Hamiltonian was constructed based on the 100ns-relaxed β-helix model, based on the same coupling models as for the 2D-IR calculations. To determine the orientation of the protein from the amide-I SSP and PPP lineshapes and intensities for 5 and 20 °C, we calculated 10,000 spectra as a function of θ and Ψ (with a grid size of 2.5°) and determined the deviation between calculation and experiment for each (θ, Ψ) point. Subsequently, Levenberg-Marquardt least-square fits were performed with the best (θ, Ψ) values as the initial guess.

## AUTHOR INFORMATION

### Author Contributions

The manuscript was written through contributions of all authors. All authors have given approval to the final version of the manuscript. ‡These authors contributed equally.

## ACKNOWLEDGMENT

This article is part of a project that has received funding from the European Research Council (ERC) under the European Union’s Horizon 2020 research and innovation programme (Grant agreement No. 819039 F-BioIce). TW, SJR and TŠT thank the Villum Foundation for financial support (Experiment Grants 22956 and 23175). KF and TŠT also acknowledge the support by The Danish National Research Foundation (Grant agreement no.: DNRF106, to the Stellar Astrophysics Centre, Aarhus University). We thank Sean A. Fischer for support with the spectral calculations, and Rolf Mertig for help with the Mathematica code used to perform the spectral fits. We thank Daniel Otzen for support with CD spectroscopy. SA would like to acknowledge funding from the National Science Foundation Graduate Research Fellowship Program under Grant No. DGE-1762114 to support this research. The simulations were done on the Hyak supercomputer system funded in part by the STF at the University of Washington. We thank Andrey Kajava and Peter Davies for providing us with structure files for the InaZ models.

