## Supporting information for "The Ice Nucleating Protein InaZ is Activated by Low Temperature"

### Experimental Details

#### Biochemistry

The InaZ9R construct was prepared essentially as described previously for a 16 repeat InaZ construct.[1] Briefly, a PCR product encoding the N-terminal domain, repeats 1-4 and 63-67 and the C-terminal domain was prepared by splicing-by-overlap PCR (SOE-PCR) using the 67 repeat *inaZ* gene from the *P. syringae* strain R10.79 as template and cloned into the pET30 Ek/LIC vector according to the manufacturer's instructions (Novagen, Merck Biosciences). This generated a recombinant construct where the truncated InaZ sequence was fused with a tag-sequence containing a His-tag and an S-tag encoded by the vector and a TEV cleavage site (introduced via the cloning primers) in the N-terminal of the construct. This construct was termed InaZ9R (see the full InaZ9R sequence in Scheme 1B and Figure S2). The InaZ9R plasmid was transformed into the *E. coli* Rosetta (DE3) strain and large-scale liquid cultures were grown in LB medium at 37$^{o}$C until an OD600 of 0.6-0.8 and induced over night at 20$^{o}$C using 1mM IPTG. The cells were lysed in lysis buffer (20 mM Tris-HCl pH 7.5; 500 mM NaCl; 20 mM Imidazole) with protease inhibitors added (cOmplete protease inhibitor cocktail, Roche). The protein was purified using immobilized metal affinity chromatography using a HisTrap FF crude column (GE Healthcare) equilibrated in lysis buffer and eluted with elution buffer (20 mM Tris-HCl pH 7.5; 500 mM NaCl; 500 mM Imidazole). Fractions containing the InaZ9R were pooled, concentrated and loaded on a MonoQ 5/50 GL column (GE Healthcare) for anion exchange chromatography. The column was equilibrated in buffer A (20 mM Tris-HCl pH 7.5; 50 mM NaCl) and eluted with a 20 column volumes of buffer B in a gradient of 0-100% (20 mM Tris-HCl pH 7.5; 1000 mM NaCl). Fractions containing the InaZ9R protein were pooled, concentrated and loaded on a Superose 6 Increase 10/300 GL column (GE Healthcare) equilibrated in SEC buffer (20 mM Tris-HCl pH 7.5; 150 mM NaCl) for size exclusion chromatography (see Figure S 1). Fractions containing InaZ9R were pooled and flash frozen in liquid nitrogen and stored at -80 °C for later use. Based on SDS-PAGE analysis (Figure S 1), the purified protein had a homogeneity of >90%.

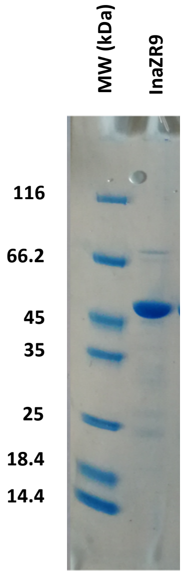

*Figure S 1:* ***SDS-PAGE of purified InaZ9R.*** *Left lane is loaded with molecular-weight marker (with the molecular weight of the marker bands shown to the left of the gel) and right lane is loaded with the peak fraction from size-exclusion column.*

**Protein sequence coverage: 56%**

Matched peptides shown in ***bold red***.

| | | | | | |
|---|---|---|---|---|---|
| 1 | MHHHHHHSSG | LVPRGSGMLE | TAAALFERQH | MDSPDLGTDD | DDKEN**LYFQG** |
| 51 | **MNLDK**ALVLR | TCAN**NMADHC** | **GLIWPASGTV** | **ESK**YWQSTRR | HENGLVGLLW |
| 101 | **GAGTSAFLSV** | **HADARWIVCE** | **VAVADIITLE** | **EPGMVK**FPR**A** | **EVVHVGDRIS** |
| 151 | **ASHFISARQA** | **DPASTPTPTP** | **TPMA**TPTPTP | AAANVALPVA | EQASHEVFDV |
| 201 | ALVSAAAPPV | NTLPVTTPQN | LQTAT**YGSTL** | **SGDNHSRLIA** | **GYGSNETAGN** |
| 251 | **HSD**LIAGYGS | TGTAGSDSSL | VAGYGSTQTA | GGDSALTAGA | DSNQTAGDR**S** |
| 301 | **KLLAGNNSYL** | **TAGDRSKLTG** | **GHDCTLMAGD** | **QSR**LTAGK**NS** | **VLTAGARSKL** |
| 351 | **IGSEGSTLSA** | **GEDSTLIFR**L | WDGKR**YRQLV** | **ARTGENGVEA** | **DIPYYVNEDD** |
| 401 | **DIVDKPDEED** | **DWIEVE** | | | |

*Figure S 2: **Sequence coverage of inaZ by tryptic and chymotryptic digest.** Purified inaZ subjected to digestion by trypsin and chymotrypsin was analyzed by mass spectrometry. Peptide sequences identified (p<0.05) are shown in bold red in the protein sequence.*

#### Mass spectral analysis of recombinant inaZ.

The recombinant inaZ9R produced was assayed by mass spectrometry to confirm protein sequence and (denatured) holo-mass of the produced protein in comparison to the nucleotide sequence of the construct. This can reveal unexpected post-translational and chemical modifications as well as amino-acid substitutions. In addition, we assayed the protein by native-spray mass spectrometry to determine possible complex formation and different folding states of the protein.

To this end, the purified protein was subjected to digest by trypsin with chymotryptic activity (owing to the limited number of fully tryptic sites in the protein) and analyzed by data dependent tandem mass spectrometry on a timsTOFpro mass spectrometer (Bruker Daltonik, Germany). Data were analyzed by fragment ion searches performed by an in-house MASCOT server (matrix science, UK.) against the sequence of inaZ alone and in the background of the *Escherichia coli* proteome database, the latter to assay the preparation for background proteins stemming from the recombinant production of the protein. Searches were conducted with cysteine carbidomethylation and methionine oxidation set as fixed and variable modifications respectively, against the inaZ sequence. This showed a maximum of 56% sequence coverage of the proposed sequence, which, owing to the limited number of fully tryptic cleavage sites, is a good coverage for this protein. In addition, the data was searched against the same inaZ sequence in the background of the *E. coli* proteome, to assay for purity of the preparation, inaZ was the major protein found in the sample (47% coverage), with a number of background proteins from *E. coli* also identified (minor components). To assay for possible unexpected modifications of inaZ, we also analyzed the data using PEAKS studio X+ (BSI bioinformatics) using its *de novo* search algorithm and found no indication of unexpected post-translational modifications to the protein (data not shown).

In addition to bottom up analysis of tryptic digests, we also assayed the intact protein by denatured and native spray mass spectrometry on a SynaptG2 mass spectrometer (Waters, UK). In short, inaZ was diluted in 50% acetonitrile, 49.9% water, 0.1% formic acid to an approximate concentration of 1 µM, and sprayed into the mass spectrometer by loading the solution into a glass-tip emitter (New Objective, USA). The mass spectrometer was externally calibrated for a range of 500-5000 m/z by sodium cesium iodide clusters and source settings were as follows: capillary voltage 1200 V, cone voltage 45 V, source gas 0.3 bar. The raw spectrum is shown in supplemental figure S3 and, although noisy, we are able to extract numerous components with a mass around 45 kDa (expected mass 43.834 kDa), which is highly similar to the mass of ~45 kDa observed on SDS page. To ascertain whether inaZ might form different folding structures with differing charge states or form complexes with itself in solution, we also attempted to measure inaZ under native conditions, using the same source settings. In this case, inaZ was diluted to approximately 1 µM in 1 mM ammonium acetate at pH 5.0. This, however, resulted in a very undefined mass spectrum (data not shown) that could not be interpreted. This could be due to the heterogeneity already shown in the poorly defined denatured spectrum being exacerbated by multiple folding or aggregation states of the protein under native conditions.

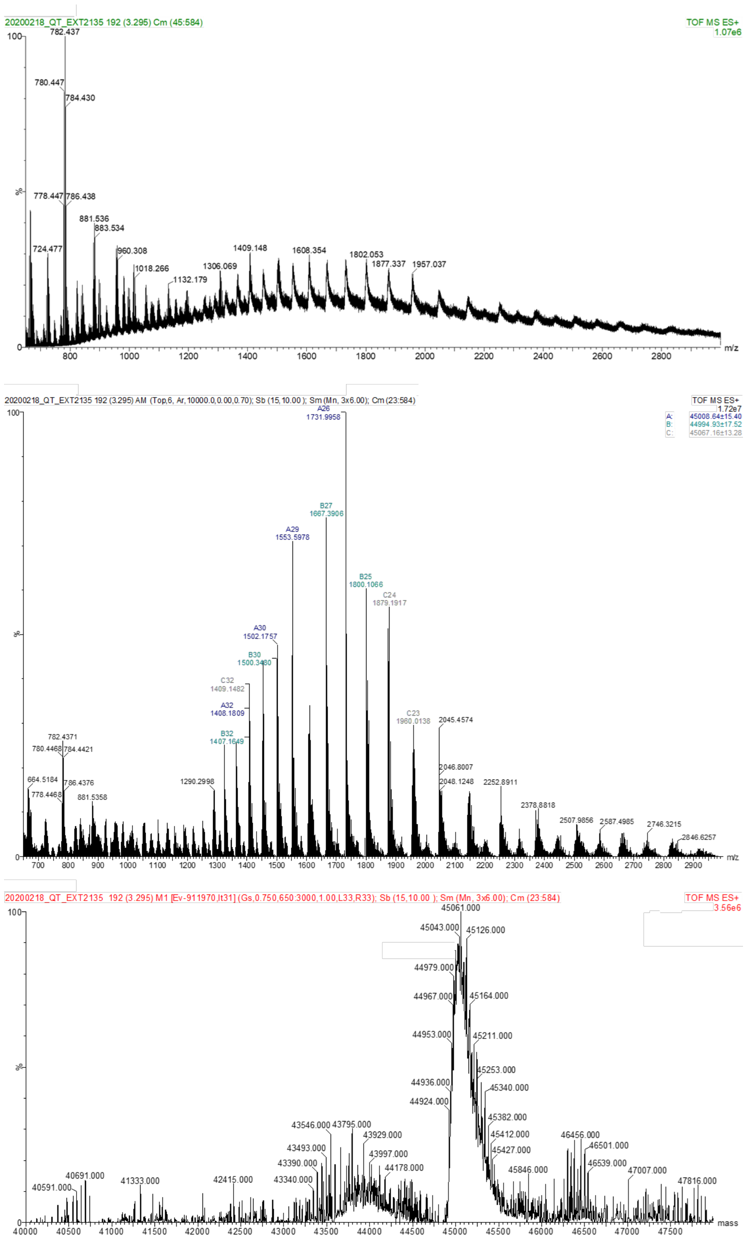

*Figure S3: **Mass spectra of intact inaZ.** The top panel shows the mass spectrum acquired from intact inaZ diluted in 50% acetonitrile, 49.9% water and 0.1% formic acid. The middle panel shows the processed spectrum (smoothed, background subtracted and centered masses) in which various components are annotated manually by the component extraction function of masslynx 3.4 (Waters, UK). Owing to the relatively noisy spectra, various components can be extracted with varying degrees of mass accuracy around a mass of 45 kDa. The bottom panel shows the automated deconvolution of the top spectrum using the MaxEnt-I algorithm of masslynx, which agrees with the manually annotated components showing a broad peak around 45 kDa with only a poorly defined component around the expected mass of 43 kDa.*

#### IR and SFG sample preparation: lyophilization

InaZ9R was lyophilized before all experiments, so that the SFG and IR amide-I spectra would not be affected by the $H_2O$ bending mode (1643 $cm^{-1}$) that overlaps with the amide-I (1600-1700 $cm^{-1}$) region. Normalized UV-CD spectra before and after lyophilization indicate that the secondary structure of the protein is not affected by this procedure (see Figure S4).

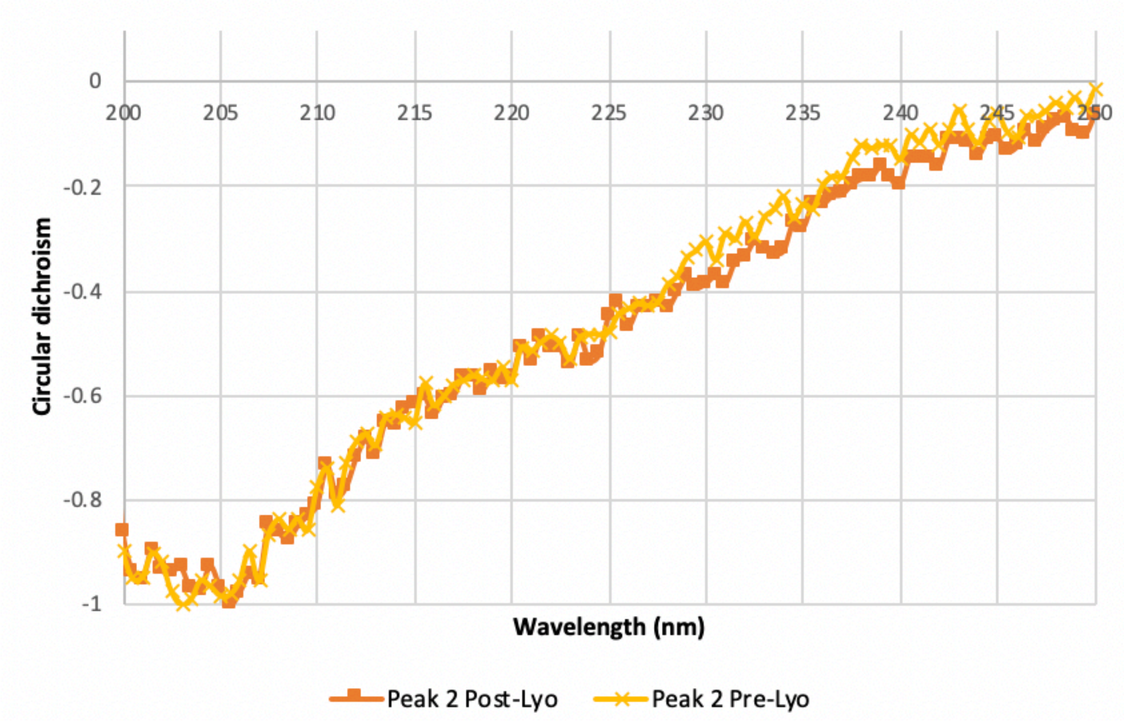

*Figure S4:* ***UV-CD spectra of InaZ9R before and after lyophilization.***

The UV-CD spectra exhibit a random coil contribution around 205 nm, as well as a broad peak around 217 nm that probably indicates beta-helical contents.[2] These circular dichroism (CD) spectra were recorded with a Chirascan-plus CD spectropolarimeter (Applied Photophysics, Leatherhead, UK), using a 1 mm pathlength quartz cuvette. The inaZ9R was dissolved in PBS buffer to a concentration of 0.05 mg/ml. The spectra were obtained at room temperature in the spectral area of 190 nm to 250 nm set to 0.5s time-per-point, step size of 0.2 nm, and 2 nm bandwidth.

#### FTIR and 2DIR Spectroscopy

The lyophilized inaZ9R in powder form was resolvated at 0.8 mg/mL in phosphate buffered saline (PBS; 0.01 M phosphate buffer, 0.0027 M KCl, and 0.137 M NaCl, pD 7.8, Sigma–Aldrich), $D_2O$ (99.9%D, Eurisotop), and 7 µL of the solution was placed in between two 1 mm $CaF_2$ windows and sealed off with a Krytox vacuum greased (Duniway) 50 µm spacer in a custom-made IR cell. The FTIR spectra were recorded after this, with 32 scans on a Bruker Vertex v70 spectrometer.

The 2DIR setup has been described before.[3] Briefly, 794 nm pulses were generated with a Mantis (Coherent) oscillator at a 80 MHz rate, of which 1 kHz were amplified with a Legend (Coherent) optical parametric amplifier to a 3 mJ beam. This light is converted to ~15 µJ of mid-IR (~6100 nm) light, with a full-width-at-half-max (fwhm) of ~150 $cm^{-1}$, after which it is split into a probe, reference and pump beam at a 5/5/95% ratio. The pump beam is spectrally narrowed to a fwhm of 10 $cm^{-1}$ by a Fabry-Pérot interferometer and overlapped in space with a 1.5 ps delay in the sample with the probe beam, while the reference beam passes through the sample cell a few mm beside the focus of the probe and pump. The difference absorbance spectrum is then dispersed by an OrielMS260i spectrograph onto a 32 pixel MCT array with a resolution of 3.9 $cm^{-1}$.

#### SFG Spectroscopy

The samples for SFG measurements were prepared from lyophilized inaZ9R powder in $D_2O$-PBS buffer to avoid interference from $H_2O$ bending modes. The samples were prepared in a stainless-steel trough with a quartz window at the bottom containing approximately 3 mL of solution. Throughout all SFG experiments, the water level was held constant by a syringe pump (New Era Pump Systems Inc.) with the cannula submerged to the bottom of the trough. A solution of $D_2O$-PBS with inaZ9R was added to the trough such that the final protein concentration was 10 $\mu$M. Throughout the experiment, a water chiller (Neslab RTE-101) was used with a water-cooled breadboard to adjust the temperature of the protein solution to 20°C, 10°C, and 5°C, as monitored with a submerged digital thermometer (Omega).

The SFG laser setup has been described previously.[4] Briefly, the setup is based on a 7 W, 35 femtosecond laser system (Astrella, Coherent) with pulses centered at 800 nm and a repetition rate of 1 kHz. One part of the output was used to pump an optical parametric amplifier (OPA) with a non-collinear difference frequency generation (NDFG) extension (TOPAS Prime, Light Conversion) to generate broadband (FWHM ~ 300 $cm^{-1}$) IR pulses tunable over the range 3.3-6.1 µm. A narrowband (FWHM ~ 15 $cm^{-1}$) visible beam was generated by guiding 1 mJ of the fundamental through a Fabry-Perot etalon. The visible and IR beams were spatially and temporally overlapped on the surface. The SFG signal was focused into a spectrograph (Shamrock 303i, Andor) and detected by an EMCCD camera (Newton 971, Andor). In the present study, SFG measurements were recorded in the amide I region (1600-1700 $cm^{-1}$), O-D stretching region (2200-2800 $cm^{-1}$), and C-H stretching region (2800-3100 $cm^{-1}$). Spectra in the Amide I region were collected in ssp (s-SFG, s-visible, p-IR) and ppp polarization combinations. Spectra in the O-D region were recorded in ssp polarization combination and spectra in the CH region were recorded in ssp and sps polarization combinations. The sample stage and IR beam path were flushed with nitrogen to avoid artifacts due to IR light adsorption by water vapor. All spectra were background subtracted and normalized using a reference spectrum obtained from gold.

#### SFG fitting methods

The SFG data in the OD region were fit using the following equation:

$$\chi_{eff}^{(2)}(\omega) = \chi_{NR}^{(2)} + \sum_q \frac{A_q}{\omega - \omega_q + i\Gamma_q} \quad (1)$$

where $\Gamma_q$, $A_q$, and $\omega_q$ are the full width half max (FWHM), amplitude, and resonant frequency of the q$^{th}$ vibrational mode, respectively, and $\chi_{NR}^{(2)}$ and $\chi_{eff}^{(2)}$ are the nonresonant background and effective seconded nonlinear susceptibility tensor, respectively. To determine the error in the measurements, the amplitude was allowed to change and the other fitting components are held the same (this is done to directly compare the amplitude change). We determine the error for the amplitudes by allowing the amplitudes to change until the fit became unreasonable. This error was determined to be 10 percent.

#### Spectral Calculation Methods

The spectra were calculated based on the amide-I Hamiltonian model.[5] Briefly, we construct a one- and two-exciton Hamiltonian for the amide-I mode of the backbone amide groups in the protein, with couplings that are estimated differently for nearest- and nonnearest-neighbor amide groups. The nearest-neighbor interactions, which are dominated by through-bond effects, are modeled using a parameterized map of an *ab initio* calculation with the 6-31G+(d) basis set and the B3LYP functional, thus providing the coupling as a function of the dihedral angle.[6, 7] The nonnearest-neighbor interactions, which are dominated by through-space effects, are estimated with the transition-dipole coupling model.[8] The local-mode IR frequencies $\nu_{local}$ are shifted with the same model employed by Lu et al.[9]

$$\nu_{local} = \epsilon_0 - \alpha * (r_{C=O} - r_{C=O\ equi.})$$

The values of $\epsilon_0$ and $\alpha$, as well as the number of frames over which to average the C=O length, was calibrated to LK$\alpha_{14}$ data and simulations using the same force field as in the current study, and found to be 1660 cm$^{-1}$, 500 cm$^{-1}$/Å and 1, respectively. The equilibrium bond distance $r_{C=O\ equi.}$ of C=O, was set to 1.229 Å for amide groups with secondary amines, which corresponds to the equilibrium value of a C-O bond in the AMBER99SB-ILDN force field, and to 1.232 Å for amide groups with tertiary amines (prolines), for which the local-mode frequency is also redshifted by 26.3 cm$^{-1}$ to account for the redshift due to the larger carbon mass as compared to the hydrogen mass bound to the amide N atom in other amino acids.

The Hamiltonian is then diagonalized to obtain the amide-I eigenvalues and eigenvectors, from which the spectroscopic response is calculated.

### Results of Spectral Calculations

#### Calculated 2DIR spectra

The effect of adding the inhomogeneous broadening to the calculated 2DIR spectra can be seen in Figure S5:

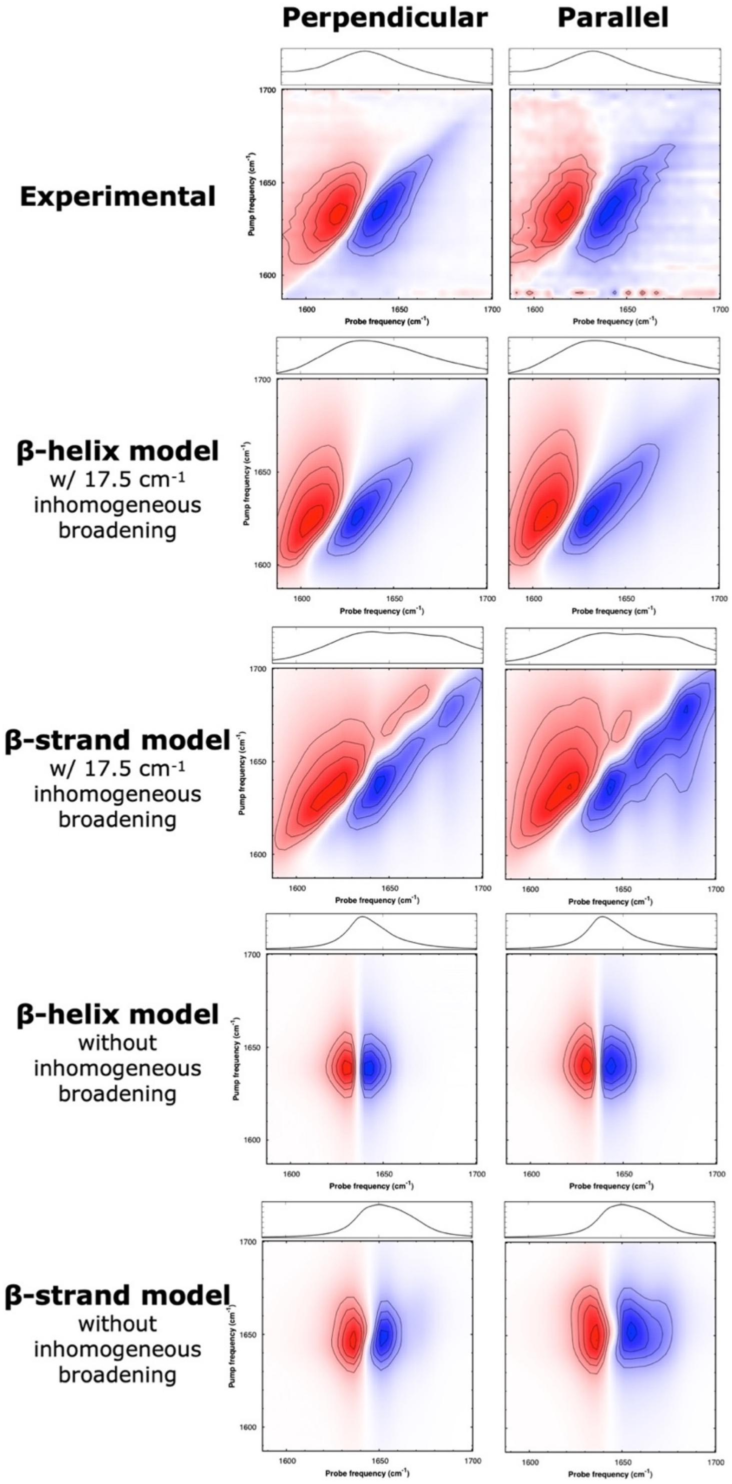

*Figure S5:* ***Comparing experimental FTIR and 2DIR spectra with several methods of spectral calculations for a parallel and perpendicular pump-probe polarization.*** *The spectra indicate that the β-helix model leads to the closest match with the experimental spectra, both with and without a random-coil contribution added to the calculations in the form of an inhomogeneous broadening of the local-mode frequencies.*

One can see that while the peak frequency is already predicted well by applying the spectral calculations to the beta-helical model, the match with the spectral shape is better with inhomogeneous broadening applied to the local-mode frequencies. In order to do this, the spectral calculation was run 50 times on the first frame of the trajectory with 50 times a gasdev[10] random distributed inhomogeneous broadening of 17.5 $cm^{-1}$ applied. The IR calculations were performed with a Lorentzian half-width-at-half-max (hwhm) of 7.5 $cm^{-1}$ and a Gaussian pump hwhm of 2.5 $cm^{-1}$. The spectra that were not inhomogeneously broadened, were calculated by averaging the hyperpolarizabilities of 250 equispaced frames of the 10 ns trajectories (thus spaced by 40 ps).

#### Calculated SFG spectra

To account for the azimutal isotropy of the proteins at the interface, we average the Euler angle $\phi$ from 0 to 2π. For the SFG spectral calculations, the total Lorentzian width was set to 20 $cm^{-1}$, in accordance with the experimentally determined visible bandwidth of 15 $cm^{-1}$ plus an inhomogeneous broadening of 5 $cm^{-1}$. Furthermore, the interfacial refractive indices were all set to 1.18 in accordance with ref. [11].

The orientational SFG investigation employed in this study requires a single (frame of a) hypothetical structure with the z-axis of the molecular frame aligned along a symmetry axis (the long axis of the β-helix in this case) so that the azimuthal averaging can be performed well in the SFG calculations. We did not include the hydrogen bond-induced shifts applied in the IR calculations (on the MD trajectories), because for a single frame this often leads to spectral distortions as not all hydrogen-bonded states are probed well in a single frame. For a good match between the calculated and experimental spectra, the gas-phase frequency was set to 1645 $cm^{-1}$, 5 $cm^{-1}$ redshifted with respect to the values used in previous studies in which similar calculations were performed[12, 13, 14, 15]. This redshift is probably necessary to account for the stronger hydrogen bonding within the β-helix as compared to the globular proteins investigated in the other studies. Both for θ and Ψ we assumed a Gaussian broadening of the orientation distribution with $\sigma$ = 10°, to account for the fact that the protein is not expected to have a very well-defined orientation distribution. To keep the spectral calculations as simple as possible, we chose to not include a lipid C=O contribution, but to focus on optimizing the match between the model and the experimental protein signal between 1600-1670 $cm^{-1}$, which is expected to be unaffected by the tail of the lipid peak (the peak centered at 1730 $cm^{-1}$).

The reported (θ,Ψ) values reported in the main text are determined by starting Levenberg-Marquardt fits with the minima of the 2D-RSS plots taken as the initial guesses. The errors are defined as the value for which the RSS doubles in a procedure in which the angles were fixed to values away from the optimal values, and performing fits in which the other fit parameters were left free. Under the azimuthal symmetry assumption and for small twist angles (θ), the uncertainty for the twist angle Ψ is relatively large, because for such angles Ψ becomes increasingly similar to the in-plane rotation angle $\phi$.

#### Additional Experimental Results

##### SFG Spectra in the C-H region

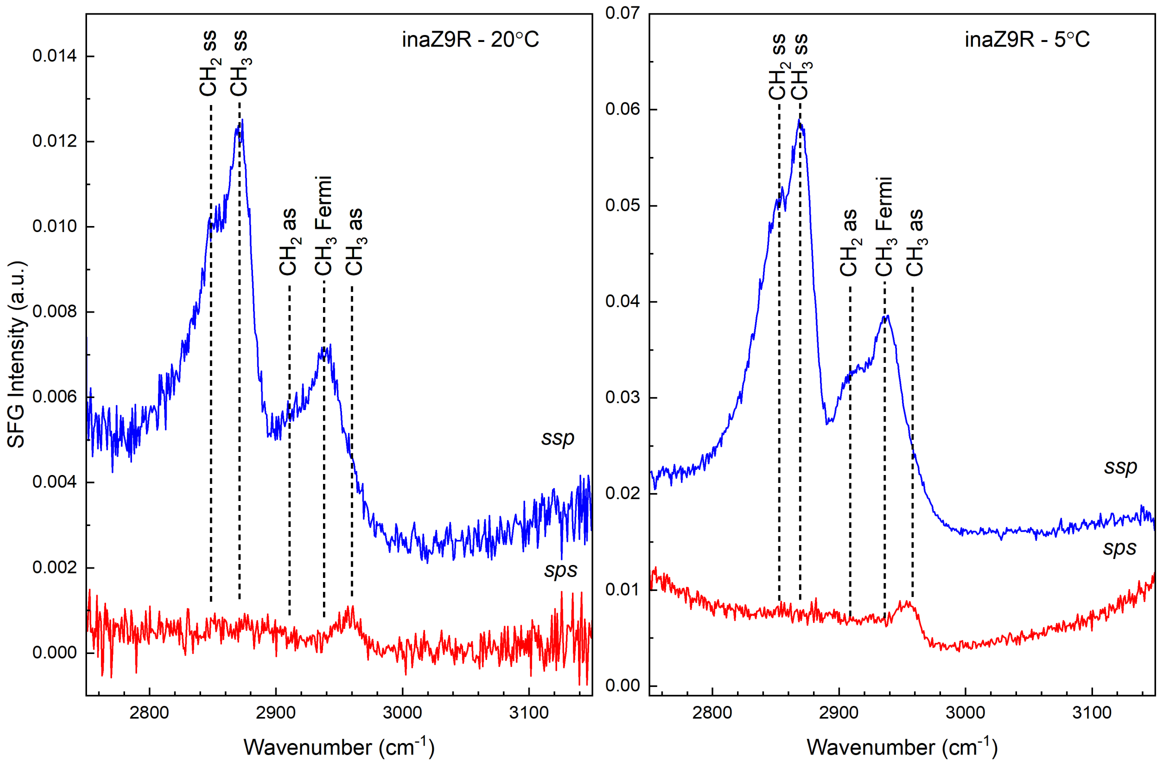

*Figure S6:* ***SFG spectra of inaZ9R at the air/$D_2O$ interface in the C-H strecth region.***

Figure S6 depicts the C–H stretch region SFG spectra of inaZ9R at the air–water interface at temperatures of 20 °C (left) and 5 °C (right). The observed C-H spectra are indicative of protein side chains and resonances are observed near 2850, 2870, 2910, 2935, and 2950 cm$^{-1}$. These are assigned to $CH_2$ symmetric, $CH_3$ symmetric, $CH_2$ asymmetric, $CH_3$ Fermi, and $CH_3$ asymmetric resonances, respectively. The observed increase in overall intensity upon decreasing the temperature from 20 °C to 5 °C, can be attributed to inaZ9R ordering or changing orientation at the interface as temperature decreases.

#### Peak Fitting Parameters

The parameters obtained from fitting the SFG data from Figure 1 of the main text are shown below in Table S1 (for the spectra with inaZ9R) and Table S2 (for the spectra without inaZ9R).

*Table S1.* ***SFG spectral parameters for the OD region:*** *air/$D_2O$ PBS interface with inaZ9R.*

| Sample/polarization | $A_{NR}$ | $\varphi_{NR}$ | $\omega_n$ (cm$^{-1}$) | $\Gamma_n$ (cm$^{-1}$) | $A_n$ | assignment |
|---|---|---|---|---|---|---|
| inaZ9R 20°C *SSP* | -0.000 | 5.53 | 2363 | 84 | 4.198 (± 0.420) | O-D stretch when a bio-molecule such as a protein or lipid is present[16] |
| | | | 2408 | 123 | 7.000 (± 0.700) | O-D stretch[16, 17] |
| | | | 2485 | 130 | 7.378 (± 0.738) | O-D stretch[17] |
| | | | 2655 | 150 | 0.411 (± 0.041) | From free O-D and protein interaction[16] |
| inaZ9R 10°C *SSP* | 0.013 | 5.53 | 2363 | 84 | 5.297 (± 0.530) | O-D stretch when a bio-molecule such as a protein or lipid is present[16] |
| | | | 2408 | 123 | 8.116 (± 0.812) | O-D stretch[16, 17] |
| | | | 2485 | 130 | 10.013 (± 1.001) | O-D stretch[17] |
| | | | 2655 | 150 | 0.532 (± 0.053) | From free O-D and protein interaction[16] |
| inaZ9R 5°C *SSP* | 0.006 | 5.53 | 2363 | 84 | 7.230 (± 0.723) | O-D stretch when a bio-molecule such as a protein or lipid is present[16] |
| | | | 2408 | 123 | 10.641 (± 1.064) | O-D stretch[16, 17] |
| | | | 2485 | 130 | 11.837 (± 1.184) | O-D stretch[17] |
| | | | 2655 | 150 | 1.371 (± 0.137) | From free O-D and protein interaction[16] |

*Table S2. **SFG spectral parameters for OD region:** air/$D_2O$ PBS interface.*

| Sample/polarization | $A_{NR}$ | $\varphi_{NR}$ | $\omega_n$ ($cm^{-1}$) | $\Gamma_n$ ($cm^{-1}$) | $A_n$ | assignment |
|---|---|---|---|---|---|---|
| $D_2O$-PBS 20℃ *SSP* | 0.041 | 6.32 | 2375 | 160 | 7.583 (± 0.758) | O-D stretch[17] |
| | | | 2460 | 105 | 1.902 (± 0.190) | O-D stretch[17] |
| | | | 2682 | 133 | 3.467 (± 0.347) | Free O-D[17] |
| $D_2O$-PBS 10℃ *SSP* | 0.062 | 6.32 | 2370 | 170 | 7.804 (± 0.780) | O-D stretch[17] |
| | | | 2480 | 140 | 3.047 (± 0.305) | O-D stretch[17] |
| | | | 2670 | 190 | 5.123 (± 0.512) | Free O-D[17] |
| $D_2O$-PBS 5℃ *SSP* | 0.031 | 6.32 | 2370 | 170 | 7.508 (± 0.751) | O-D stretch[17] |
| | | | 2480 | 140 | 3.586 (± 0.359) | O-D stretch[17] |
| | | | 2670 | 190 | 5.439 (± 0.544) | Free O-D[17] |

### Molecular Dynamics Simulations

#### MD Methods

The two protein structures used in this study were obtained from Andrey Kajava and Peter Davies, respectively. The stacked antiparallel β-sheet model was first simulated in a 4.7 x 5.1 x 5.7 $nm^3$ water box containing 4337 water molecules and 3 sodium counterions. The β-helix model was simulated in a 6.4 x 5.2 x 4.2 $nm^3$ water box containing 4066 water molecules and 8 sodium counterions. All simulations were performed using GROMACS 2019.[18] using the GROMOS-53A6 forcefield[19] in combination with the SPC/Ewater model.[20] First each system underwent a steepest descent energy minimization to remove any unfavorable contacts, with a tolerance of 1000 kJ $mol^{-1}$ $nm^{-1}$. Following this, the systems were equilibrated to 300 K (26.85 °C) using a stochastic velocity-rescaling thermostat[21] ($\tau$ = 0.1 ps) and to 1 bar using a Parrinello-Rahman barostat[22] ($\tau$ = 1.0 ps) over 100 ps. In all simulations, periodic boundaries were applied in all directions. All simulations were run using a timestep of 2 fs, and hydrogen bonds were constrained in all simulations by the LINCS algorithm.[23] Electrostatic interactions were calculated with the particle mesh Ewald (PME) summation method.[24] A van der Waals cutoff value of 1.0 nm was used.

Following equilibration, production simulations were carried out in solution in the NVT ensemble at two different temperatures 300K (26.85 °C), and 278K (5 °C) using the same parameters used for equilibration. The solution production run was carried out for 10 ns to assess the stability of the proposed structures in solution at both temperatures. A root-mean square deviation of the alpha-carbon atoms is included in 7. For the production simulations at the air-water interface (AWI), we ran MD of the protein in a restrained and unrestrained form in the NVT ensemble. First, the protein was restrained to the PDB predicted structure using a harmonic restraint (1000 kJ/mol) on the α-carbon atoms using the RMSD collective variable in PLUMED 2.4.2.[25]

#### MD Snapshots

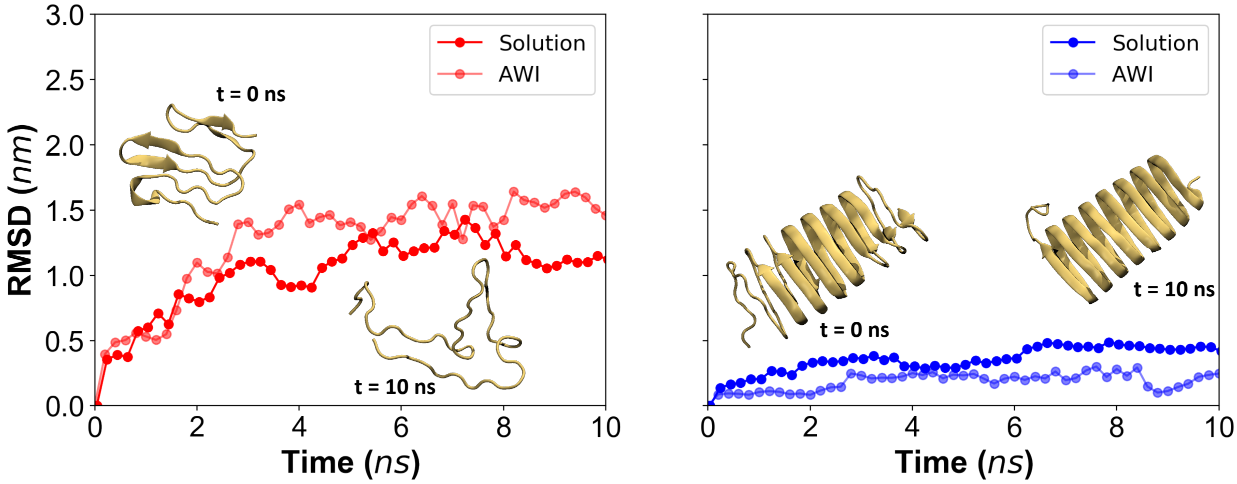

*Figure S7:* ***Stability of (Left) stacked antiparallel $\beta$-sheet and (Right) $\beta$-helix model monitored by taking the RMSD of the α-carbon in reference to frame 0.***
